# Three-dimensional imaging reveals preserved intrinsic contractile function in aging human skeletal muscle fibers

**DOI:** 10.64898/2026.06.09.730973

**Authors:** Carlos S. Zepeda, Laura E. Teigen, Isabell Dobrzycki, Yuan Wen, Christopher W. Sundberg

**Affiliations:** Department of Physical Therapy, Marquette University, Milwaukee, WI, USA; Division of Geriatrics and Gerontology, Department of Medicine, University of Wisconsin–Madison, Madison, WI, USA; University of Kentucky Center for Muscle Biology, Lexington, KY, USA; Department of Physiology, University of Kentucky, Lexington, KY, USA; Division of Biomedical Informatics, Department of Internal Medicine, University of Kentucky, Lexington, KY, USA; Athletic and Human Performance Research Center, Marquette University, Milwaukee, WI, USA; Department of Kinesiology, University of Wisconsin–Madison, Madison, WI, USA

## Abstract

Age-related reductions in muscle fiber size and contractile function, particularly in fibers expressing fast myosin heavy chains, contribute to declines in whole-muscle power. However, methodological limitations in estimating fiber size during contractile experiments have likely contributed to conflicting findings regarding whether reduced single-fiber force and power in older adults reflects their smaller size and/or impaired intrinsic contractile function. To address this, we coupled single-fiber contractile experiments with 3D-imaging in 7 young (19-40yrs) and 6 older (69-84yrs) males to assess intrinsic contractile function and compare agreement between 3D-derived cross-sectional area (CSA) and CSA estimates obtained either in air or solution. Fast fiber CSA from older males were ∼28–45% smaller across measurement conditions compared with young, whereas slow fiber CSA did not differ. Accordingly, absolute force and power of fast fibers were 41% and 37% lower. When normalized to CSA from measurements in air or 3D-imaging, size-specific force and power either did not differ or were greater in older adults, indicating preserved intrinsic contractile function in both fiber types. This was supported by no age-related differences in the rate of tension redevelopment (k_tr_), a size-independent measure of intrinsic contractile function. In contrast, size-specific force and power calculated using solution-based CSA estimates were lower in older compared with young adults, and Bland-Altman analyses demonstrated the poorest agreement between solution-based and 3D CSA measurements. These findings indicate that intrinsic contractile function is preserved with aging and suggest that methodological differences in CSA measurement contributes to the disparate findings in the literature.

## INTRODUCTION

Aging is associated with a progressive decline in skeletal muscle mass and function (strength and power) that leads to reduced mobility and quality of life in older adults. Notably, the age-related decline in muscle power (∼2-4% yr^-1^) occurs more rapidly than the loss of total muscle mass (∼0.5-1% yr^-1^) (Mitchell *et al*., 2012; Alcazar *et al*., 2020; Alcazar *et al*., 2023), suggesting that factors beyond muscle atrophy, such as impairments in the ability of the nervous system to activate the muscle and/or factors disrupting intrinsic contractile function, must also contribute to the loss in power. Importantly, the age-related decline in the ability to generate power is a stronger predictor of physical impairments (Reid & Fielding, 2012) and mortality (Araujo *et al*., 2025) than muscle strength. Despite the importance of muscle power to mobility and physical function in older adults, the mechanisms for the accelerated age-related loss in power relative to muscle mass are not well understood.

It was recently reported that the reduced ability of older adults to generate power in the lower limb is due primarily to factors within the muscle, with neural factors potentially playing a minor role in older females (Wrucke *et al*., 2024). However, identifying the mechanisms within the muscle remains particularly challenging, in part due to the fiber type heterogeneity and the well-established differential effects of aging on the size and function of each fiber type (Lexell *et al*., 1988; Nilwik *et al*., 2013; Teigen *et al*., 2020; Grosicki *et al*., 2022; Teigen *et al*., 2026). One proposed mechanism underlying the accelerated age-related decline in whole muscle power is the selective atrophy of the fibers expressing the fast myosin heavy chain (MyHC) isoforms, which often exhibit marked reductions in force- and power-generating capacity with aging (Sundberg *et al*., 2018a; Sundberg *et al*., 2025). Indeed, a recent review of the aging single fiber literature found that the fast fibers from older adults are typically smaller and generate lower absolute force and power compared with young adults, whereas slow fiber size and contractile function are largely preserved (Grosicki *et al*., 2022). However, when fast fiber force and power were normalized to fiber size, which is the most commonly reported metric of intrinsic contractile function, the findings are inconsistent, with studies reporting impaired intrinsic contractile function in older adults (Larsson *et al*., 1997; Frontera *et al*., 2000; D’Antona *et al*., 2003; Ochala *et al*., 2007; Hvid *et al*., 2013; Lamboley *et al*., 2015; Brocca *et al*., 2017), no differences (Korhonen *et al*., 2006; Yu *et al*., 2007; Raue *et al*., 2009; Harber *et al*., 2012; Teigen *et al*., 2020; Teigen *et al*., 2026), or even enhancements with aging (Straight *et al*., 2018; Sundberg *et al*., 2018a; Gries *et al*., 2019; Grosicki *et al*., 2021; Sundberg *et al*., 2025). Thus, despite nearly three decades of research, it remains unknown whether the reductions in fast fiber force and power in older adults are attributed to their smaller size and/or impaired intrinsic contractile function.

One potential critical, yet underappreciated, factor contributing to this longstanding inconsistency may be a fundamental methodological limitation in how muscle fiber cross-sectional area (CSA) is estimated (Smith & Herzog, 2023; Mebrahtu *et al*., 2024; Hessel *et al*., 2026). In chemically permeabilized and mechanically peeled fiber preparations, CSA is estimated from two-dimensional (2D) images by measuring the fiber diameter at multiple points while the fiber is either submerged in relaxing solution or briefly suspended in air (< 5s). The use of air-based measurements arose from the recognition that fibers in solution are not simple geometric shapes (Blinks, 1965; Malakoutian *et al*., 2021), but instead exhibit irregular cross-sectional profiles that violate the conventional assumptions of circular or elliptical fiber geometry. Suspending fibers in air was therefore intended to leverage surface tension and capillary forces to induce a more uniform, cylindrical shape, thereby better approximating these geometric assumptions (Allen & Moss, 1987; Metzger & Moss, 1987). However, even under these conditions, it is unknown how well these measurements approximate the fiber CSA, and variations in the exposure time to air can differentially affect the fiber dimensions (Ferenczi, 1986), particularly in fibers from older adults (unpublished observations). Furthermore, studies assessing muscle fiber shape using immunohistochemical methods have reported that fibers from older adults exhibit greater deviations from circularity, particularly the fast MyHC II fibers (Kirkeby & Garbarsch, 2000; Barnouin *et al*., 2017; Horwath *et al*., 2024; Soendenbroe *et al*., 2024). Collectively, these limitations suggest that inaccuracies in CSA estimation from 2D images may contribute to the inconsistent findings on intrinsic contractile function with aging.

Continued advancements in cellular imaging and computational analyses, particularly using confocal microscopy, have enabled high-resolution, three-dimensional (3D) renderings of isolated human muscle fibers (Liu *et al*., 2009; Cristea *et al*., 2010), allowing more precise measurements of fiber morphology. However, no study has combined single-fiber contractile mechanics with 3D morphological analysis of the same fiber segments or directly compared 3D-derived CSA with conventional CSA estimates derived from 2D diameter measurements in solution and air. Thus, the primary aims of this study were to: 1) determine whether intrinsic contractile function is impaired with aging by assessing single fiber contractile mechanics followed by 3D morphological analysis of the same fibers, and 2) compare the agreement between CSA estimates obtained from 2D images in solution and air with those derived from 3D imaging. We hypothesized that 1) reductions in fast fiber force and power with aging will be determined primarily by their smaller size rather than altered cross-bridge mechanics, with no age differences observed in slow fibers; 2) CSA estimates from 2D images obtained in both solution and air would show bias and poor agreement with 3D-derived measurements; and 3) methodological differences in fiber CSA assessment would account for much of the discrepancies in the literature regarding whether aging alters intrinsic single-fiber contractile function.

## METHODS

### Participants and ethical approval

Seven young (20 – 40 yr) and 6 older males (69 – 84 yr) volunteered and provided written informed consent to participate in this study. Participants underwent a general health screening and were excluded from participation if they were taking medications that affect muscle mass or neuromuscular function (e.g. hormone-replacement therapies, antidepressants, glucocorticoids, etc.). Participants were healthy, community-dwelling adults free of any known neurological, musculoskeletal, and cardiovascular diseases. All experimental procedures were approved by the Marquette University Institutional Review Board and conformed to the principles in the Declaration of Helsinki.

### Physical activity assessment

Physical activity for each participant was quantified using a triaxial accelerometer (GT3X; ActiGraph, Pensacola, FL, USA) worn around the waist for at least 4 days (2 weekdays and 2 weekend days) as reported previously (Hassanlouei *et al*., 2017; Sundberg *et al*., 2018a). The data were recorded for each participant if the accelerometer was worn for a minimum of 8 hours on at least 3 days (Hart *et al*., 2011).

### Experimental overview

Participants reported to the laboratory on two occasions, once for a muscle biopsy and again to measure thigh lean mass and whole-muscle function of the knee extensor muscles. The purpose of the whole muscle session was to determine whether older adults exhibited typical age-related differences in knee extensor mass and function, including lower thigh lean mass as well as lower absolute and mass-specific mechanical force and power outputs.

### Thigh lean mass

Body composition and thigh lean mass were assessed with dual X-ray absorptiometry (Lunar iDXA; GE, Madison, WI, USA). Thigh lean mass was quantified for the region of interest using the manufacturer’s software (enCORE 14.10.022; GE), with the distal demarcation set at the tibiofemoral joint and the proximal demarcation set as a diagonal bifurcation through the femoral neck. DXA measures of thigh lean mass with these landmarks are strongly correlated with measures from magnetic resonance imaging but underestimate the age-related loss in thigh muscle mass (Maden-Wilkinson *et al*., 2013).

### Whole-muscle knee extensor function

Whole-muscle function of the knee extensors was measured using the same experimental setup described previously (Sundberg *et al*., 2018b). Briefly, participants were seated upright in the high Fowler’s position on a Biodex System 4 dynamometer (Biodex Medical, Shirley, NY, USA) with the dominant leg (preferred kicking leg) positioned at 90° knee flexion. Participants were secured using the dynamometer’s four-point restraint system to minimize extraneous movements and changes in hip angle, and they were prohibited from grasping the dynamometer with their hands to ensure the measured torques and velocities were generated primarily by the knee extensor muscles.

Following a standardized warmup consisting of dynamic knee extension exercise lifting a light load (1 Nm), participants performed a minimum of three brief (2-3 s) maximal voluntary isometric contractions (MVCs) separated by a minimum of 60 s rest. Strong verbal encouragement and visual feedback were provided via a 56 cm monitor mounted 1-1.5 m directly in front of the participant. MVC torque was quantified as the average torque over a 0.5 s interval centered on peak torque, and MVC attempts were continued until the two highest values were within 5% of each other. Participants then completed 4 MVCs combined with electrical stimulation to assess voluntary activation and involuntary contractile properties as described previously (Delgadillo *et al*., 2021).

### Muscle stimulation and electrically evoked contractile properties

Involuntary contractile properties of the knee extensors were assessed using transcutaneous electrical stimulation delivered through custom pad electrodes (6 × 8 cm), with the cathode positioned approximately 8–12 cm proximal to the superior border of the patella and the anode was positioned 10–13 cm proximal to the cathode. Stimulations were delivered using a constant-current stimulator (DS7AH, Digitimer, Hertfordshire, UK), and the stimulator intensity established using single-pulse stimuli (400 V, 200 μs duration) beginning at 50 mA and increasing incrementally by 50–100 mA until the amplitude of the unpotentiated resting twitch torque plateaued. Stimulus intensity was then increased by an additional 20% to ensure supramaximal stimulation. This intensity was subsequently used for all paired-pulse (100 Hz doublet) stimulations during the voluntary activation and contractile property assessments.

Participants performed MVCs during which a 100 Hz paired-pulse stimulus was delivered both during the contraction and immediately following the MVC (< 5 s). Each MVC trial was separated by at least 2 min of rest. The reported values for potentiated resting twitch amplitude (Nm), half-relaxation time (ms), and peak rate of torque development (Nm·s^-1^) were calculated as the median values from the four MVC trials. The peak rate of torque development was quantified as the derivative of the torque signal as the highest rate of torque increase over a 10 ms interval.

Following the isometric MVC measurements, participants were habituated to performing maximal velocity knee extensions against a 20% MVC load applied by the dynamometer. Participants were instructed to kick as fast as possible once every 3 s for a total of 4 min (80 contractions). Peak power was the highest power output recorded from the first 10 contractions (30 s) of the task. Peak power obtained from this single-load fatiguing protocol has previously demonstrated strong associations and agreement with values derived from a comprehensive torque–velocity assessment, supporting its validity for evaluating age-related differences in muscle power (Wrucke *et al*., 2026).

### Voluntary activation

Voluntary activation of the knee extensors was assessed using the interpolated twitch technique (Herbert & Gandevia, 1999). During the four MVCs, transcutaneous muscle stimulation was applied and voluntary activation calculated with Equation 1:

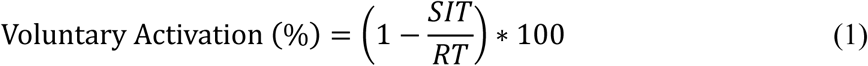

where SIT is the amplitude of the superimposed twitch torque elicited by the stimulation during the MVC and RT is the electrically-evoked, potentiated resting twitch immediately after the MVC (< 5 s). The reported voluntary activation for each participant was the median obtained from the 4 MVC trials.

### Muscle biopsy

A muscle biopsy from the *vastus lateralis* was obtained from each participant using the modified Bergström technique as described previously (Bergstrom, 1975; Sundberg *et al*., 2018a). Participants were instructed to abstain from strenuous exercise of the lower limbs for 48 h prior to the biopsy and arrive at the laboratory fasted and without the consumption of caffeine for ≥ 8 h. Longitudinal bundles from the biopsy sample were immediately placed in ice cold glycerol skinning solution (see below) and stored at −20 °C for up to 4 weeks. The remaining bundles of the biopsy sample were orientated longitudinally on a small notecard, flash frozen in liquid nitrogen-cooled isopentane and stored at −80 °C until sectioned for immunohistochemistry (IHC).

### Single fiber morphology and contractile mechanics experiments

*Solutions:* The composition of the relaxing (pCa 9.0, where pCa = -log[Ca^2+^]) and activating (pCa 4.5) solutions used for the single fiber contractile experiments were derived from an iterative program using the stability constants adjusted for temperature, pH, and ionic strength (Fabiato & Fabiato, 1979; Fabiato, 1988). The solutions contained (in mM): 20 imidazole, 7 EGTA, 4 MgATP and 14.5 creatine phosphate. Inorganic phosphate (P_i_) was added as K_2_HPO_4_ to yield a concentration of 4 mM. The ionic strength was adjusted to 180 mM with KCl, and the pH adjusted to 7.0 with KOH and HCl. The skinning solution was composed of 50% relaxing solution and 50% glycerol (vol:vol).

### Experimental setup

Single fiber segments ∼5-10 mm in length were isolated from the biopsy sample and prepared as described previously (Sundberg *et al*., 2018a). Briefly, the ends of the fiber were secured with 4.0 monofilament posts tied with 10.0 nylon sutures to a force transducer (400A; Aurora Scientific, Aurora, Ontario, CA) and a high-speed servomotor (controller 312B; Aurora Scientific) with ∼2-3 mm of the fiber between the two attachment points. Fibers were suspended in 100 μl of relaxing solution kept at 16 °C with a temperature-controlled Peltier unit for the duration of the experiment, except when transferred into air for imaging or to a second Peltier unit for activation at 20 °C. To view the fiber at 800x magnification, the microsystem was transferred to the stage of an inverted microscope. The sarcomere length was adjusted to 2.5 μm using a calibrated eyepiece micrometer, and the fiber length measured as the distance between the two attachment points via a mechanical micrometer (Starrett; Athol, MA, USA).

To assess differences in single-fiber cross-sectional area (CSA) measured in solution and following varying durations of air exposure, digital images were obtained while fibers were suspended in relaxing solution and while exposed to air for ≤ 2 s (air ASAP), 5 s (air 5s), and 10 s (air 10s). Fiber diameter was determined from 3 measurements obtained along a 280-μm segment centered at the exact midpoint of the fiber, as calculated from total fiber length. The average fiber diameter was then used to calculate CSA under the assumption that the fiber is a cylindrical shape.

### Force-velocity and k_tr_

Force-velocity and force-power curves were obtained as described previously (Sundberg *et al*., 2018a; Sundberg *et al*., 2025). Fibers were maximally activated in saturating Ca^2+^ (pCa 4.5), allowed to generate peak isometric force (P_o_), and then subjected to three predetermined submaximal isotonic loads (300-FC1 Force Controller; Positron Development, Inglewood, CA, USA). Fibers were activated four to six times to obtain 12-18 different isotonic loads. Each force-velocity curve was fit with an iterative non-linear curve fitting procedure (Levenberg-Marquardt algorithm) using the hyperbolic Hill equation (Hill, 1938) (Equation 2):

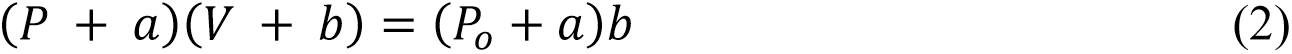

where *P* is force, *V* is velocity, *P_o_* is peak isometric force, and *a* and *b* are constants with dimensions of force and velocity, respectively (Widrick *et al*., 1996). Absolute (μN**·**fl**·**s^-1^) and size-specific power (W**·**L^-1^) were calculated as the product of shortening velocity (fl**·**s^-1^) and absolute (μN) and size-specific force (kN**·**m^-2^), respectively, and the peak fiber power determined using the fitted parameters from the force-velocity curve (Widrick *et al*., 1996). The reported absolute and size-specific P_o_ was the average of all the contractions within each condition.

Following the force-velocity experiments, the low-to high-force transition of the cross-bridge cycle was measured by quantifying the rate of force redevelopment, *k*_tr_, in response to a rapid slack re-extension maneuver of a maximally Ca^2+^-activated fiber (Metzger *et al*., 1989; Sundberg *et al*., 2018a). Only fibers producing > 90% of initial force underwent *k*_tr_ analysis. Force redevelopment following re-extension was fit with a first-order exponential function, where *k*_tr_ is the rate constant of force redevelopment (s^-1^) (Metzger *et al*., 1989), which is independent of fiber size and thus provides an additional measure of intrinsic contractile function.

### Single fiber fluorescent staining

Following single fiber contractile experiments, fibers were carefully transferred and the ends secured with 10.0 nylon sutures between two small-diameter (800 µm) stainless-steel rods within a custom-built imaging microchamber (Fig. 1). Each rod was fixed to maintain fiber alignment between the attachment points but could be adjusted in two planes to control sarcomere length and fiber height. Due to the mounting procedure on the single fiber contractile apparatus (Moss, 1979), both ends of the fiber segments became crimped at the point of attachment, creating visible landmarks that were used to ensure consistent positioning of the same fiber segment within the 3D imaging microchamber for subsequent imaging. Fibers were then stored overnight in relaxing solution at 4°C.

**Fig. 1.**
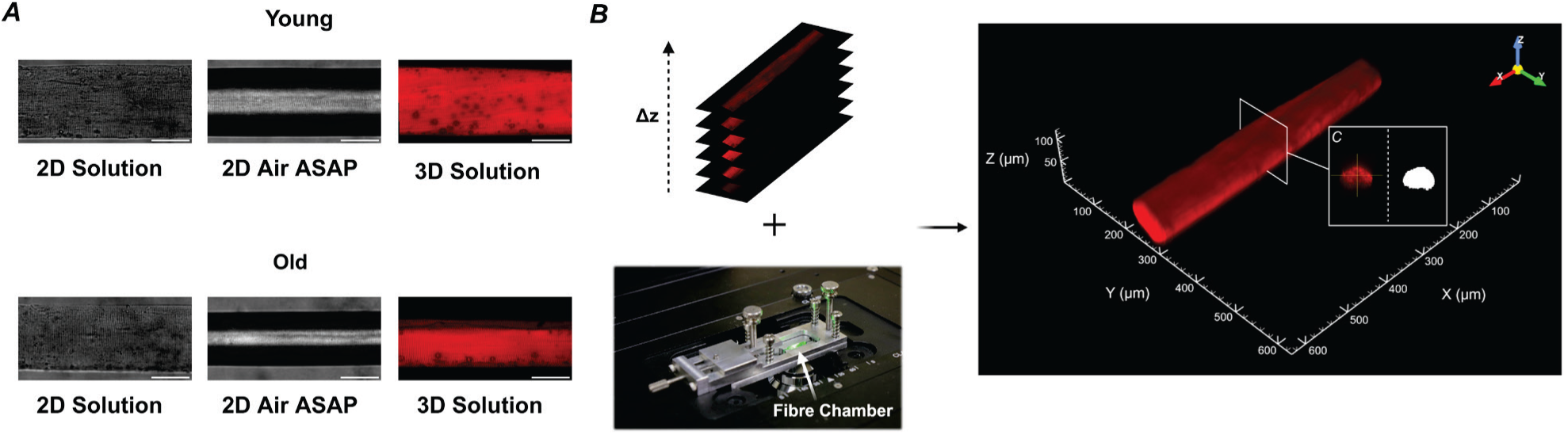
Representative images of an isolated muscle fiber obtained in air, solution, and by 3D imaging. Images of isolated muscle fibers from young and older males were acquired while the fibers were suspended in relaxing solution, in air following transfer from solution (air ASAP), or in relaxing solution with confocal microscopy for 3D reconstruction (A). Confocal z-stacks were acquired at 1-μm intervals to generate 3D renderings of isolated muscle fibers (B). Representative orthogonal slice view (left) and corresponding segmentations generated using the custom image analysis software (right) were used to measure CSA at 0.38 μm^2^ resolution across the length of the fiber.

On the day of staining and imaging, fibers were first incubated for 15 min in 0.1% Triton X-100 (T9284; Sigma-Aldrich) diluted in relaxing solution at room temperature (RT) to facilitate penetration of the fluorescent probe. After a 5 min wash in relaxing solution, fibers were incubated for 60 min in Alexa Fluor 568 phalloidin (1:1000, A12830; Invitrogen) diluted in relaxing solution at RT. Fibers were then washed 3×5 min in relaxing solution before sarcomere length was adjusted for imaging.

### Image acquisition

Prior to imaging, the microchamber was placed on the stage of an inverted microscope equipped with a calibrated eyepiece micrometer, and the sarcomere length was adjusted to 2.5 μm (as described above). Single fiber images were acquired using an inverted laser-scanning confocal microscope (Ti2-E, Nikon) equipped with 405- and 561-nm laser lines and a Plan-Apochromat 20x/0.75 NA objective lens. For 3D renderings of each fiber, consecutive images were acquired through the Z-plane at 1-μm increments (Fig. 1). The length of each fiber segment captured was approximately 635 μm. Images were acquired at 1024 x 1024 pixel resolution, yielding a total of 1024 CSA measurements along the length of each fiber. Image analysis was conducted with a custom-automated imaging analysis software developed in MATLAB (version: 9.14.0 (R2023a), The MathWorks, Natick, MA) with CSA for each fiber reported as the average CSA from all 1024 measurements. This approach reduced potential investigator bias and enabled quantification of fiber CSA at a spatial resolution of 0.38 μm^2^.

### Myosin heavy chain (MyHC) composition

MyHC composition of the isolated fibers was determined by SDS-PAGE. Following image acquisition, fibers were solubilized in 80 μL of SDS sample buffer and gels were made up of a 3% acrylamide/bis (19:1) stacking layer and 5% separating layer. Gels were stored at 4 °C for at least 24 hrs before loading and were run at 100V for ∼24 hrs at 4 °C with 0.07% β-mercaptoethanol in the top running buffer. The gels were then stained with Sypro Ruby Protein Gel Stain (S12000; Invitrogen) according to the manufacturer’s instructions and imaged using a UVP GelStudio touch gel imager (849-97-0942-01; analytikjena) with a 605nm bandpass emission filter to visualize the MyHC bands.

### Immunohistochemistry

To complement the single fiber CSA data from the contractile mechanics experiments, which are known to be limited by sample size and unintentional selection bias against the smaller fibers (Grosicki *et al*., 2022), we performed immunohistochemistry (IHC) on tissue cross-sections as described previously (Teigen *et al*., 2026). Briefly, 7 μm cross-sections were cut and stored at - 80 °C until staining for MyHC isoforms. On the day of staining, slides were rehydrated in phosphate-buffered saline (PBS) and incubated with a primary antibody (1°Ab) cocktail consisting of rabbit (Rb) α-laminin IgG (1:100, L9393, Sigma-Aldrich, RRID:AB_477163), mouse (Ms) α-MyHC I IgG2b (1:100, BA-D5-c; DSHB, RRID:AB_2235587), Ms α-MyHC IIa IgG1 (1:500, SC-71-s; DSHB, RRID:AB_2147165), Ms α-MyHC IIx IgM (1:50, 6H1-s; DSHB, RRID:AB_1157897), and PBS for 90 min at RT. Muscle sections were then washed with fresh PBS for 3×5min at RT and subsequently incubated in a secondary antibody (2°Ab) cocktail of Goat (Gt) α-Rb IgG (H+L), AMCA (1:50, CI-1000-1.5, Vector Laboratories, RRID:AB_2336195), Gt α-Ms IgG2b Alexa Fluor (AF) 647 (1:250, A21242, Invitrogen, RRID:AB_2535811), Gt α-Ms IgG1 AF488 (1:250, A21121, Invitrogen, RRID:AB_2535764), Gt α-Ms IgM AF555 (1:500, A21426, Invitrogen, RRID:AB_2535847), diluted in PBS for 60 min at RT. Sections were again washed in fresh PBS, mounted with coverslips using 1:1 PBS/glycerol, and then stored at 4 °C until imaging.

### Image acquisition

Muscle section images were acquired using a high-resolution fluorescence microscope (BZ-X810, Keyence) with a Plan Apochromat 20x mounted objective (BZ-PA20, Keyence) and filters DAPI (49000, Chroma Technology), Cy5 (49006, Chroma Technology), FITC/Alexa Fluor 488/Fluo3/Oregon Green (49011, Chroma Technology), and CY3/TRITC (49004, Chroma Technology). Automated detection software, MyoVision (Wen *et al*., 2018), was used for fiber type-specific CSA with manual verification of all fibers.

### Statistical analysis

Participant anthropometrics, whole-muscle knee extensor function, physical activity levels, and MyHC content from IHC were compared between age groups (young and old) using an unpaired *t* test. Simple linear regression analyses were performed between neuromuscular measurements (voluntary activation and electrically evoked contractile properties) and power production to examine the extent to which neural and muscular factors are associated with the age-related loss of power. Statistical analysis for anthropometrics, whole-muscle knee extensor function, physical activity levels, and IHC data were performed with SPSS, version 31.0 (IBM Corp. Armonk, NY, USA).

To test for differences in single fiber morphology and contractile mechanics between young and old males, a repeated measures mixed effects nested ANOVA was used with age group (young and old) and condition (air ASAP, air 5s, air 10s, solution, 3D) included as fixed factors. When a significant main effect of condition was observed, post hoc pairwise comparisons were performed using Tukey’s method. When a significant age x condition interaction was detected, separate mixed effects nested ANOVAs were performed within each condition to compare age groups. This was done to allow direct comparison of our findings with those from the field that use a single CSA measurement. Simple linear regression analyses were performed using each subject’s whole-muscle peak power and mean single-fiber power to examine associations between power production at the cellular and whole-muscle levels. Bland-Altman plots were used to assess mean bias and limits of agreement (LoA) between CSA estimated from 2D images (both solution and air) and CSA measured from 3D imaging. Statistical analyses for single fiber data were performed using Minitab version 22.4.0 (Minitab Inc., State College, PA, USA). Statistical significance was set at *P* < 0.05. Data are presented as the mean ± SD in the text and tables and the mean ± SE in the figures.

A total of 400 fibers were attempted, of which 21 were excluded because of experimental failure (11 failed quality-control criteria, 6 were lost during transfer to the imaging chamber, and 4 did not yield acceptable 3D images). Of the remaining 379 fibers, 2 were MyHC I/IIa hybrid fibers and were excluded, resulting in a final sample of 377 fibers. Because the number of hybrid IIa/IIx and pure IIx fibers were small and similar between age groups (young: n = 17; old: n = 23), we grouped these fibers with the pure IIa fibers and refer to the grouped data as MyHC II fibers.

## RESULTS

### Participant anthropometrics and whole-muscle knee extensor function

Anthropometrics, physical activity levels, and whole-muscle knee extensor function are presented in Table 1. Physical activity levels did not differ between young and older adults (*P* = 0.973). Older adults exhibited characteristic signs of aging skeletal muscle, including ∼37% lower thigh lean mass (*P* = 0.014), ∼50% lower absolute isometric torque (*P* < 0.001), and ∼56% lower absolute mechanical power output compared with young adults (P < 0.001, Fig. 2A). After normalizing to thigh lean mass, mass-specific isometric torque (*P* = 0.014) and power (*P* = 0.012) remained 22% and 31% lower in older adults.

**Fig. 2.**
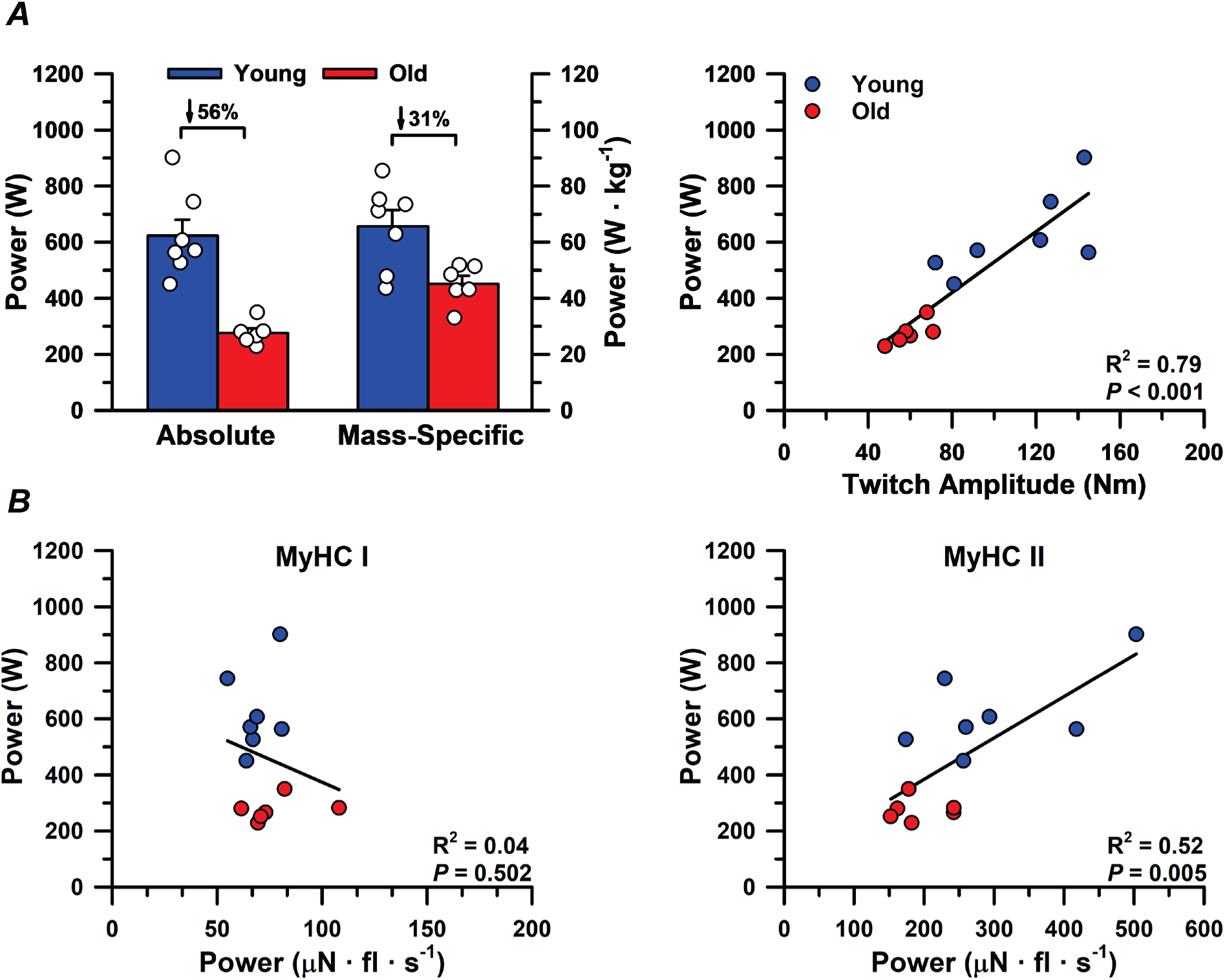
Whole-muscle knee extensor power output in young and older males. Mean absolute peak power output was 56% lower in the older compared with young adults and remained 31% lower after normalization to thigh lean mass (A). Linear regression analysis revealed that absolute peak power was strongly associated with the amplitude of the potentiated twitch (A), suggesting that intramuscular factors are a major contributor to age-related differences in whole-muscle power. Absolute whole-muscle power was associated with the mean power output of fast myosin heavy chain (MyHC) II fibers, but not slow MyHC I fibers (B), suggesting that age-related reductions in fast fiber output is an important factor contributing to differences in whole-muscle power.

### Voluntary activation, electrically evoked contractile properties, and associations with whole-muscle power

Voluntary activation did not differ between young (87.3 ± 7.4%) and older adults (89.3 ± 7.6%; *P* = 0.631). Resting potentiated twitch amplitude (young: 111.7 ± 29.8 Nm, old: 60.0 ± 8.5 Nm) and the rate of twitch torque development (young: 1086.7 ± 300.7, old: 573.3 ± 100.9 Nm**·**s^-1^) were greater in young compared with older adults (both *P* = 0.002), whereas the half relaxation time did not differ between groups (young: 83.6 ± 25.8 ms, old: 109.7 ± 25.4 ms; *P* = 0.094).

Simple linear regression analyses revealed that several muscular factors were moderately to strongly associated with whole-muscle peak power, including thigh lean mass (R^2^ = 0.57; *P* = 0.003), twitch amplitude (R^2^ = 0.79; *P* < 0.001, Fig. 2A), half relaxation time (R^2^ = 0.48; *P* = 0.008), and the rate of twitch torque development (R^2^ = 0.77; *P* < 0.001). In contrast, voluntary activation was not associated with peak power (R^2^ = 0.00; *P* = 0.876). Linear regression analyses examining the relationship between peak whole-muscle power and mean single-fiber absolute power from each subject revealed no association for MyHC I fibers (R^2^ = 0.04; *P* = 0.502). However, fast MyHC II single-fiber absolute power was associated with whole-muscle absolute power (R^2^ = 0.52; *P* = 0.005, Fig. 2B). After normalizing to whole-muscle and single-fiber size, mass-specific power was not associated with size-specific power for either MyHC I (R^2^ = 0.04; *P* = 0.506) or MyHC II fibers (R^2^ = 0.05; *P* = 0.477).

### Agreement between estimated fiber CSA from 2D images and 3D measurements

Bland-Altman plots were used to assess agreement between CSA estimated from 2D images obtained in air and solution and CSA measured from 3D imaging in 377 fibers pooled across age groups and fiber types (Fig. 3). The smallest mean bias was observed for CSA estimated after 10 s of air exposure (mean bias = 136 μm^2^; LoA = −2,387 to 2,657 μm^2^) (Fig. 3C), indicating the closest agreement with 3D measurements. Larger biases were evident for air 5s (mean bias = 1,064 μm^2^; LoA = −1,602 to 3,732 μm^2^) and air ASAP (mean bias = 1,977 μm^2^; LoA = −1,060 to 5,013 μm^2^) (Fig. 3A,B), whereas the greatest bias was observed with CSA estimated in solution (mean bias = 3,609 μm^2^; LoA = −2,866 to 10084 μm^2^) (Fig. 3D). Regression analyses revealed moderate-to-strong associations between CSA estimated from the 2D imaging conditions and 3D CSA measurement (R^2^ = 0.53-0.78; *P* < 0.001) (Fig. 3). Notably, a vast majority of the data points fell below the line of identity (x = y), indicating systematic overestimation of CSA compared with 3D CSA measurements, consistent with the positive mean bias observed in each condition.

**Fig. 3.**
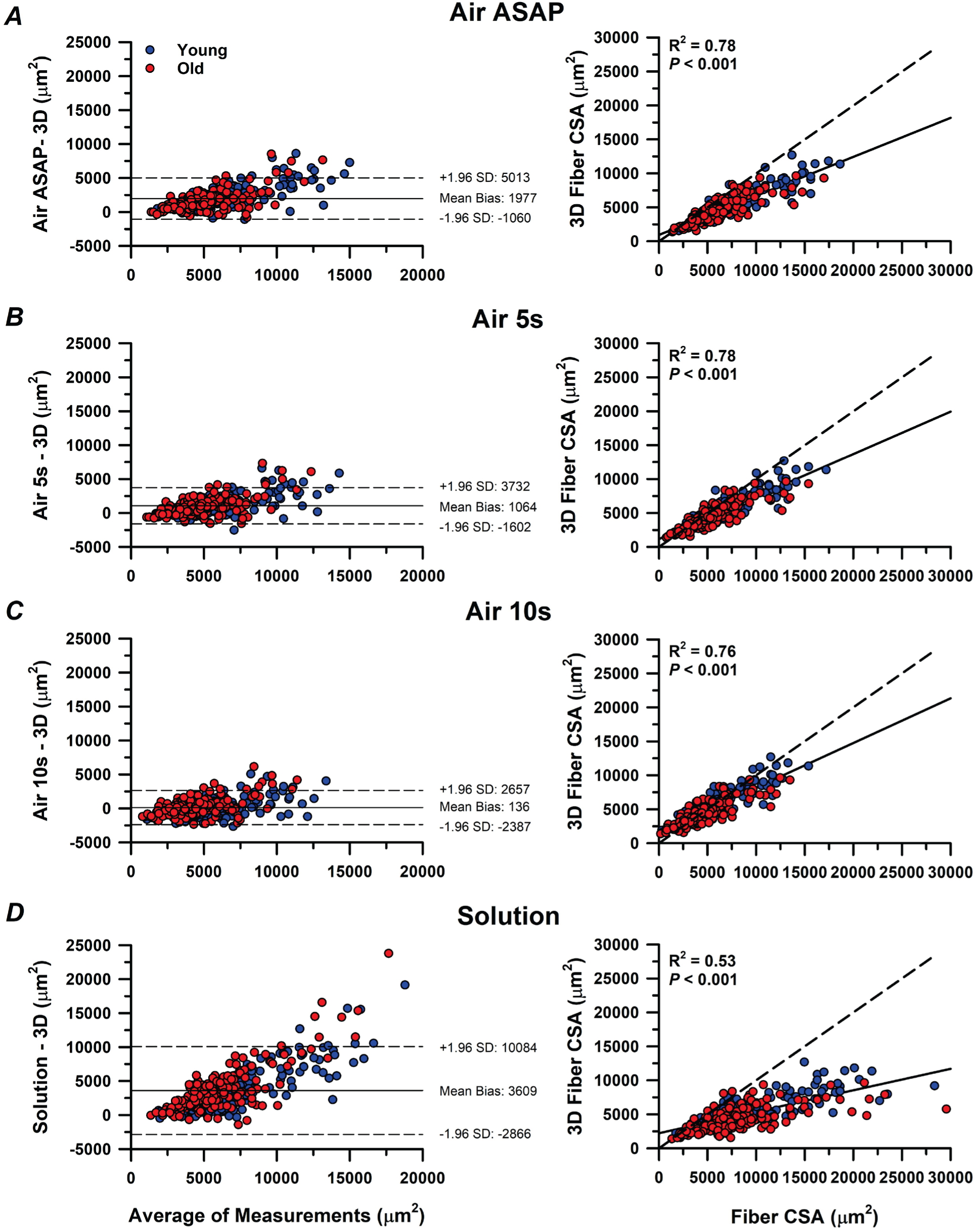
Bland-Altman plots illustrating the mean bias and limits of agreement (± 1.96 SD) between CSA estimates obtained from 2D images and CSA derived from 3D imaging. CSA estimates from 2D images were obtained in air as soon as possible following transfer from solution (ASAP; A), after 5 s in air (B), after 10 s in air (C), and while suspended in relaxing solution (D). A total of 377 muscle fibers (Young = 210; Old = 167), including both fiber types were pooled across age groups. In the Bland-Altman plots (left), the dotted lines represent the LoA and the solid line represents the mean bias. In the regression plots (right), the dotted line represents the line of equality (x = y), and the solid line represents the fitted regression line. Each data point represents an individual fiber.

### Single muscle fiber morphology and contractile mechanics

#### Slow MyHC I fibers

CSA measured across conditions for the 177 MyHC I fibers (young = 97, old = 80), along with the corresponding IHC data, are presented in Fig. 4 and Table A1. There was no effect of age when all CSA measurements were combined (*P* = 0.279), but there was an age x condition interaction (*P* = 0.001). However, separate mixed effects nested ANOVA analyses revealed no differences in MyHC I CSA between young and older adults in air ASAP (*P* = 0.662), air 5s (*P* = 0.394), air 10s (*P* = 0.281), solution (*P* = 0.076), or 3D (*P* = 0.514). Independent of age, there was an effect of condition (*P* < 0.001), with CSA estimated from 2D images in solution being 22-66% larger than all other conditions (*P* ≤ 0.001 for all). A clear hierarchy in CSA measurement was observed such that CSA in solution > air ASAP > air 5s > 3D > air 10s (*P* ≤ 0.008 for all).

**Fig. 4.**
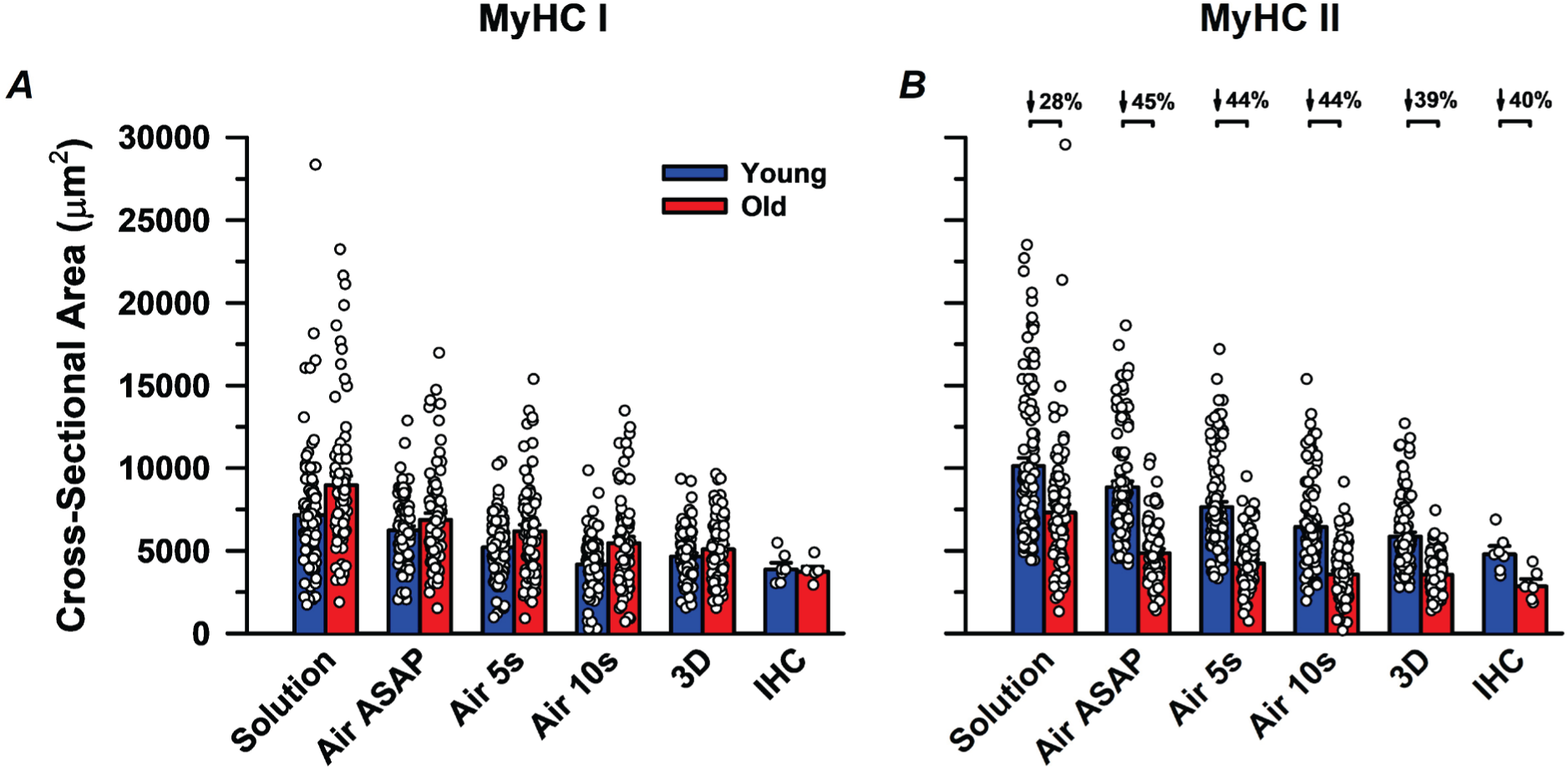
Cross-sectional area (CSA) of single muscle fibers estimated using different measurement approaches (Solution, Air ASAP, Air 5s, Air 10s, and 3D imaging) and via immunohistochemistry (IHC). Myosin heavy chain (MyHC) I fiber CSA did not differ with age across any single-fiber measurement approach or when assessed via IHC (A). In contrast, MyHC II fiber CSA was markedly smaller in older compared with young males across all single-fiber measurement approaches and via IHC (B). Significance was set at *P* < 0.05. Data points represent individual fibers for the single-fiber CSA measurements and participant means for the IHC analyses. Values are presented as mean ± SEM.

There was no effect of age on the percent reduction in fiber CSA measured in air relative to solution (*P* = 0.132); however, there was an effect of condition (*P* < 0.001) and an age x condition interaction (*P* < 0.001). Specifically, MyHC I fibers from older adults had a greater reduction in CSA from solution to air ASAP (19 ± 21%) compared with young adults (5 ± 19%; *P* = 0.021), but no age-related differences were observed for air 5s (old: 29 ± 18%, young: 22 ± 17%, *P* = 0.124) or air 10s (old: 38 ± 18%, young: 38 ± 17%, *P* = 0.936). Independent of age, the percent reduction in CSA relative to solution was greatest in air 10s, followed by air 5s and air ASAP (*P* < 0.001 for all).

Mean force-velocity curves, absolute P_o_, and maximal shortening velocity (V_max_) for MyHC I fibers are presented in Fig. 5 and Table A1. There were no differences between young and older adults in the curvature of the force-velocity relationship (a/P_o_, *P* = 0.531), absolute P_o_ (*P* = 0.341), or V_max_ (*P* = 0.745). In addition, the rate of force redevelopment (k_tr_) did not differ between young (8.36 ± 0.79 s^-1^) and older adults (7.74 ± 1.28 s^-1^, *P* = 0.349).

**Fig. 5.**
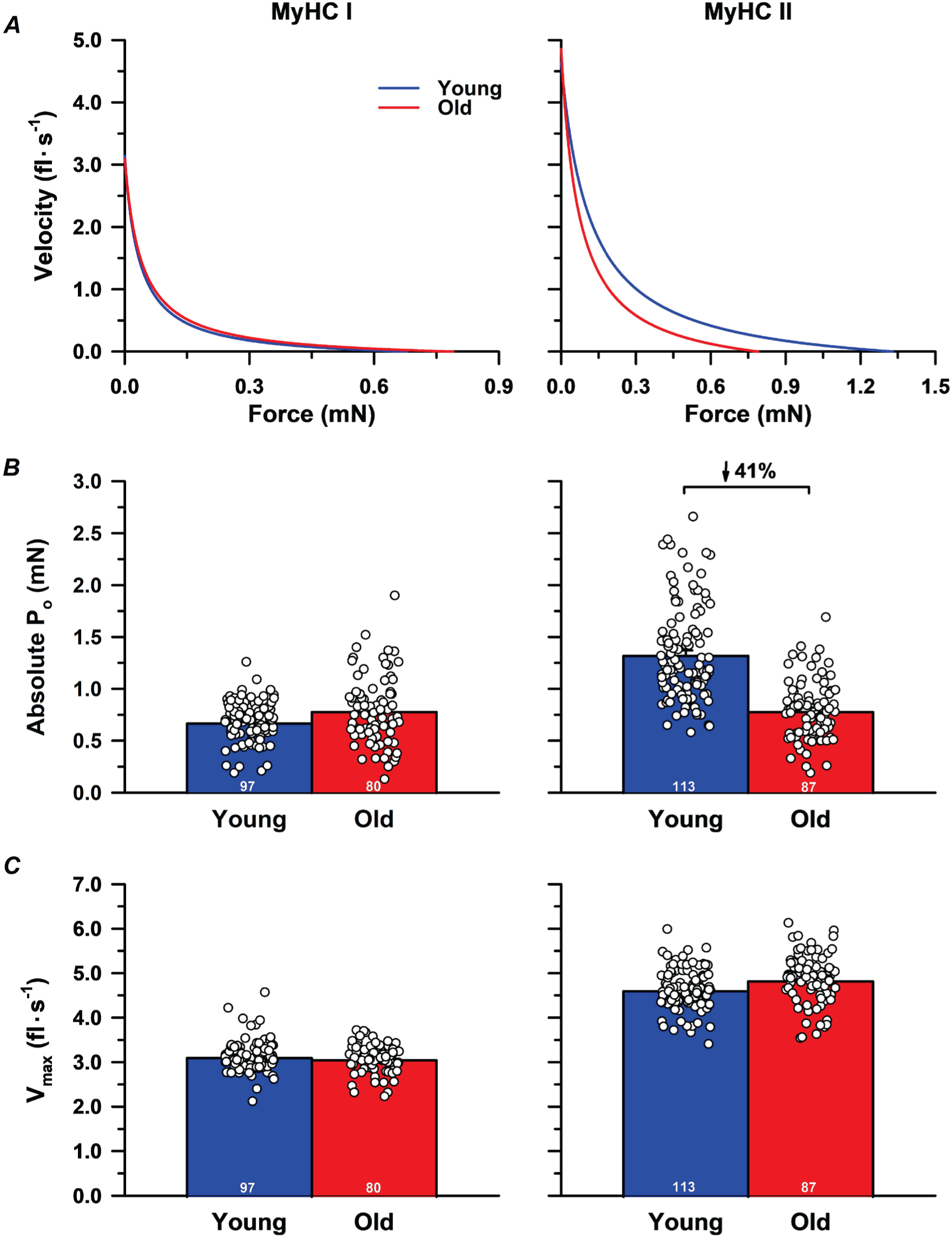
Force-velocity curves, peak isometric force (P_o_), and maximal shortening velocity (V_max_) of slow myosin heavy chain (MyHC) I and fast MyHC II fibers. The curvature of the force–velocity relationship (a/P_o_) did not differ between young and older males in either fiber type (A). Absolute P_o_ did not differ with age in MyHC I fibers but was 41% lower in MyHC II fibers from older compared with young males (B). V_max_ did not differ between young and older males in either fiber type (C). Significance was set at *P* < 0.05. Each data point in the bar graphs represents an individual fiber, with the number of fibers (*n*) displayed within the bars. Values are presented as mean ± SEM.

Size-specific and absolute force-power curves for slow MyHC I fibers are presented in Fig. 6 and Table A1. There was no effect of age on size-specific P_o_ when all CSA measurements were combined (*P* = 0.449), but there was an age x condition interaction (*P* < 0.001). However, separate mixed effects nested ANOVA analyses revealed no differences in size-specific P_o_ between young and older adults in any condition (air ASAP, *P* = 0.370; air 5s, *P* = 0.667; air 10s, *P* = 0.176; solution, *P* = 0.088; 3D, *P* = 0.541). Independent of age, there was an effect of condition (*P* < 0.001), where size-specific P_o_ was lowest when normalized to CSA estimated from 2D images in solution compared with all other conditions (*P* ≤ 0.001 for all, Fig. 6). A clear hierarchy of CSA normalization was observed such that size-specific P_o_ air 10s > 3D > air 5s > air ASAP > solution (*P* < 0.001 for all).

**Fig. 6.**
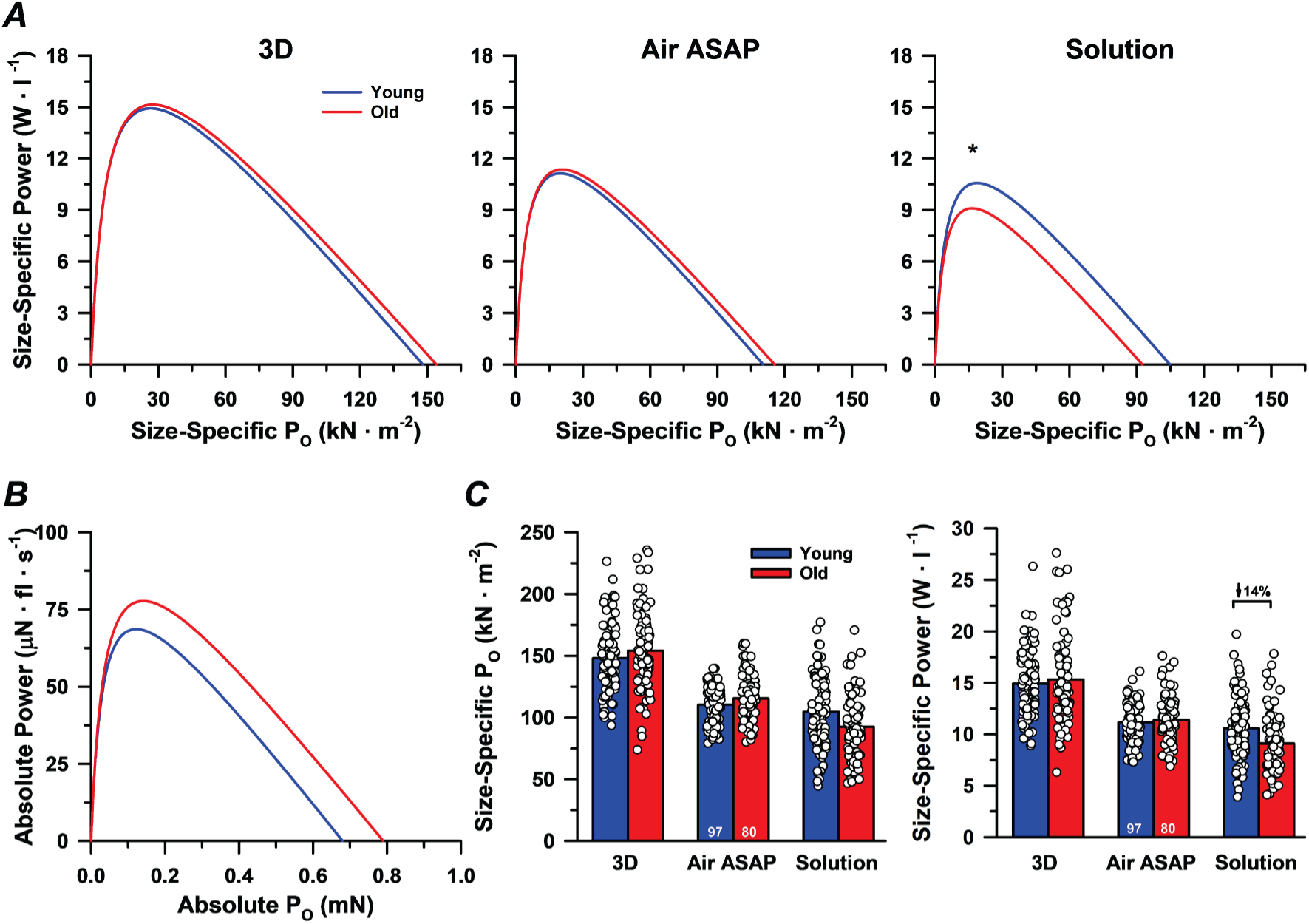
Force-power curves and size-specific force and power of slow myosin heavy chain (MyHC) I fibers. Size-specific force (P_o_) of MyHC I fibers did not differ between young and older males regardless of the CSA measurement approach used (A). Similarly, absolute P_o_ and power output did not differ with age (B). Size-specific power output calculated with 3D-derived CSA or air ASAP measurements did not differ with age; however, size-specific power output was 14% lower in older compared with young males when calculated with solution-based CSA estimates (A, C). Data from the air 5s and air 10s conditions were omitted, as air ASAP is the only air-based measurement approach reported in previous single-fiber studies. Significance was set at *P* < 0.05. Each data point in the bar graphs represents an individual fiber, with the number of fibers (*n*) displayed within the bars. Values are presented as mean ± SEM.

Absolute peak power of slow MyHC I fibers did not differ between young and older adults (*P* = 0.238). There was also no effect of age on size-specific peak power when all CSA measurements were combined (*P* = 0.292), but there was an age x condition interaction (*P* < 0.001). Specifically, size-specific peak power was ∼14% lower in older (9.1 ± 2.8 W·L^-1^) compared with young from the 2D images in solution only (10.6 ± 2.9 W·L^-1^, *P* = 0.033), whereas no age-related differences were observed in air ASAP (*P* = 0.900), air 5s (*P* = 0.415), air 10s (*P* = 0.112), or 3D (*P* = 0.614). Independent of age, size-specific peak power was greatest when CSA was measured after 10 s of air exposure compared with all other conditions (*P* < 0.001 for all). A clear hierarchy of CSA normalization was observed such that size-specific peak power air 10s > 3D > air 5s > air ASAP > solution (*P* < 0.001 for all).

#### Fast MyHC II fibers

CSA measured across conditions for the 200 MyHC II fibers (young = 113, old = 87), along with the corresponding IHC data, are presented in Fig. 4 and Table A2. There was an effect of age (*P* = 0.003) and age x condition interaction (*P* < 0.001), where MyHC II fiber CSA was smaller in the older compared with young adults in all conditions (air ASAP, *P* = 0.002; air 5s, *P* = 0.004; air 10s, *P* = 0.005; solution, *P* = 0.034; 3D, *P* = 0.001). Consistent with the data from MyHC I fibers, there was an effect of condition (*P* < 0.001), with CSA estimated from 2D images in solution being 25-82% larger than all other conditions (*P* < 0.001 for all). Independent of age, there was a clear hierarchy in CSA measurement such that CSA in solution > air ASAP > air 5s > 3D ≈ air 10s (*P* < 0.001 for all, except 3D vs. air 10s, where *P* = 0.500).

In contrast to MyHC I fibers, there was an effect of age on the percent reduction in fiber CSA measured in air relative to solution (*P* = 0.002), along with an effect of condition (*P* < 0.001) and an age x condition interaction (*P* = 0.004). Specifically, MyHC II fibers from older adults had a greater reduction in CSA from solution to air ASAP (old: 26 ± 23%, young: 7 ± 22%, *P* = 0.007), air 5s (old: 36 ± 20%, young: 20 ± 19%, *P* = 0.005), and air 10s (old: 48 ± 18%, young: 33 ± 16%, *P* = 0.002). Similar to MyHC I fibers, the percent reduction in CSA relative to solution was greatest in air 10s, followed by air 5s and air ASAP (*P* < 0.001 for all).

Mean force-velocity curves, absolute P_o_, and maximal shortening velocity (V_max_) for MyHC II fibers are presented in Fig. 5 and Table A2. The curvature of the force-velocity relationship (a/P_o_, *P* = 0.069) and V_max_ (*P* = 0.056) did not differ between young and older adults. However, absolute P_o_ was ∼41% lower in older (0.79 ± 0.28 mN) compared with young adults (1.33 ± 0.46 mN, *P* = 0.003). In addition, the rate of force redevelopment (k_tr_) did not differ between young (18.16 ± 3.92 s^-1^) and older adults (19.20 ± 3.97 s^-1^, *P* = 0.409).

Size-specific and absolute force-power curves for fast MyHC II fibers are presented in Fig. 7 and Table A2. There was no effect of age on size-specific P_o_ when all CSA measurements were combined (*P* = 0.692), but there was an age x condition interaction (*P* < 0.001). Specifically, size-specific P_o_ was ∼15% lower in the older (119.3 ± 37.4 kN·m^-2^) compared with young from the 2D images in solution only (140.1 ± 36.7 kN·m^-2^, *P* = 0.005), whereas no age-related differences were observed in air ASAP (*P* = 0.177), air 5s (*P* = 0.158), air 10s (*P* = 0.137), or 3D (*P* = 0.667). Independent of age, there was an effect of condition (*P* < 0.001), where size-specific P_o_ was greatest when CSA was measured in air 10s and 3D compared with all other conditions (3D ≈ air 10s > air 5s > air ASAP > solution, *P* < 0.001 for all, except 3D vs. air 10s, where *P* = 1.000).

**Fig. 7.**
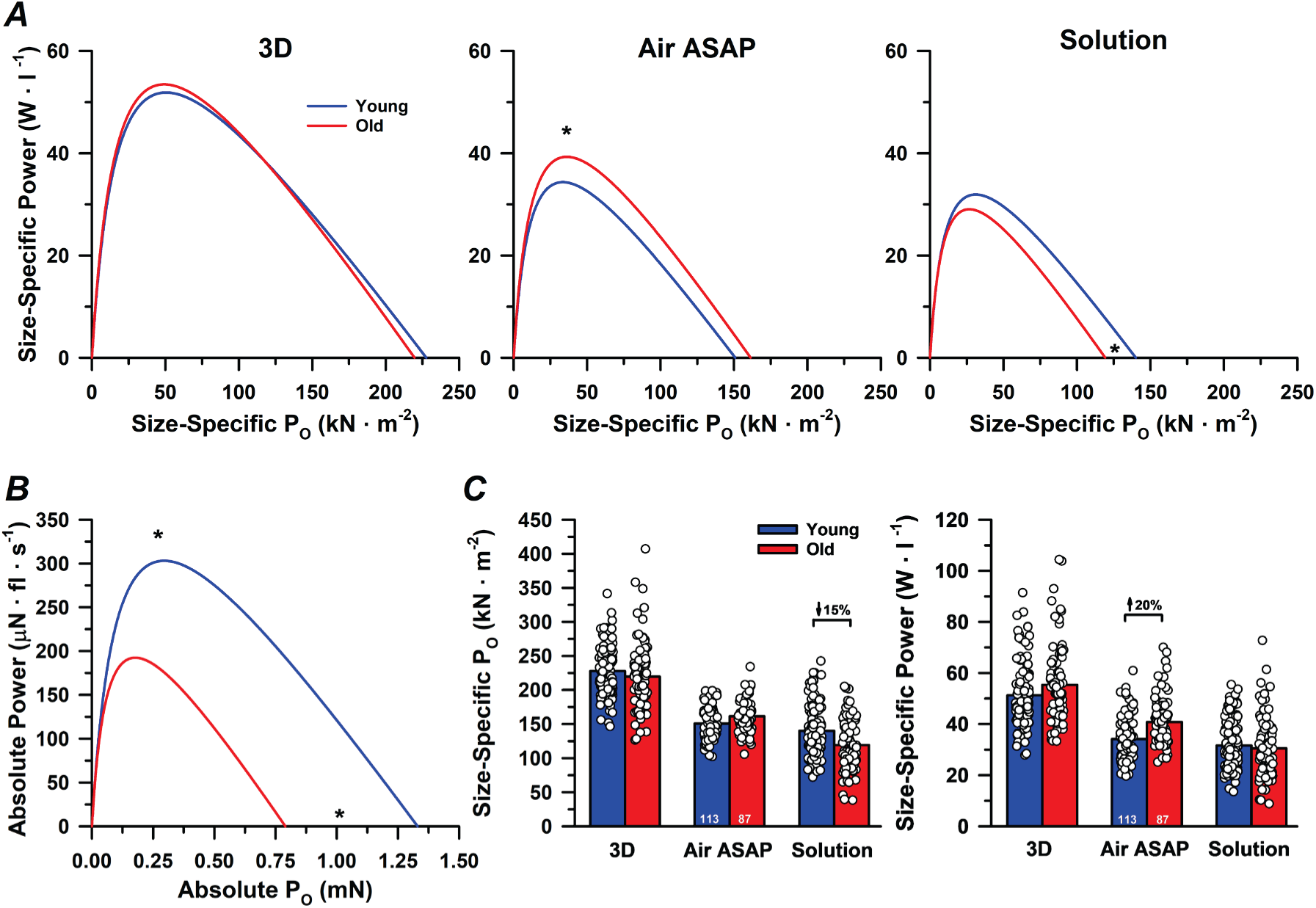
Force-power curves and size-specific force and power of fast myosin heavy chain (MyHC) II fibers. Size-specific force (P_o_) of MyHC II fibers did not differ between young and older males when normalized with 3D-derived CSA or air ASAP measurements but was 15% lower in older adults when normalized with solution-based CSA estimates (A, C). In contrast, absolute P_o_ and power output were markedly lower in older compared with young males (B). Size-specific power output calculated with 3D-derived CSA or solution-based CSA estimates did not differ with age, whereas size-specific power output calculated with air ASAP measurements was 20% higher in older compared with young males (A, C). Data from the air 5s and air 10s conditions were omitted, as air ASAP is the only air-based measurement approach reported in previous single-fiber studies. Significance was set at *P* < 0.05. Each data point in the bar graphs represents an individual fiber, with the number of fibers (*n*) displayed within the bars. Values are presented as mean ± SEM.

Absolute peak power of fast MyHC II fibers was ∼37% lower in older (192.4 ± 68.0 uN·fl·s^-1^) compared with young adults (303.2 ± 134.0 uN·fl·s^-1^, *P* = 0.020). There was also an effect of age on size-specific peak power when all CSA measurements were combined (*P* = 0.026) and an age x condition interaction (*P* < 0.001). Specifically, size-specific peak power was greater in older compared with young adults in air ASAP (*P* = 0.014), air 5s (*P* = 0.014), and in air 10s (*P* = 0.015), whereas no age-related differences were observed in solution (*P* = 0.626) or 3D (*P* = 0.346, Fig 7 and Table A2). Independent of age, there was an effect of condition on size-specific peak power (*P* < 0.001) where size-specific power was greatest when CSA was measured in air 10s and 3D compared with all other conditions (air 10s ≈ 3D > air 5s > air ASAP > solution, *P* < 0.001 for all, except air 10s vs 3D where *P* = 0.298).

### Fiber morphology and MyHC distribution from immunohistochemistry

Fiber type-specific CSA determined from 445 ± 155 fibers in the young (range: 222 – 687 fibers) and 528 ± 173 fibers in older adults (range: 256 – 801 fibers) are presented in Fig. 3. Consistent with the isolated single fiber data, MyHC I fiber CSA did not differ between age groups (young: 3,946 ± 884 μm^2^, old: 3,798 ± 628 μm^2^; *P* = 0.738), whereas MyHC II fiber CSA was ∼40% smaller in older (2,910 ± 938 μm^2^) compared with young adults (4,857 ± 1,120 μm^2^; *P* = 0.006). Accordingly, the MyHC II/I CSA ratio was greater in young (1.25 ± 0.26) compared with older adults (0.76 ± 0.16; *P* = 0.002).

Fiber type distribution did not differ between young and older adults for MyHC I (young: 50.4 ± 16.0%, old: 64.9 ± 18.8%; *P* = 0.160), MyHC IIa (young: 38.7 ± 16.4%, old: 28.0 ± 11.4%; P = 0.206), MyHC IIa/IIx (young: 9.4 ± 13.1%, old: 5.2 ± 7.6%; P = 0.506), MyHC IIx (young: 1.3 ± 3.3%, old: 1.5 ± 2.7%; P = 0.930), or combined MyHC II fibers (young: 49.4 ± 15.9%, old: 34.6 ± 18.8%, *P* = 0.153). In contrast, MyHC I proportional area was greater in older (71.4 ± 15.8%) compared with young adults (47.1 ± 16.3%; *P* = 0.020), and MyHC II proportional area was lower in older adults (28.3 ± 15.8% vs. 52.6 ± 16.3%; *P* = 0.020).

## DISCUSSION

The present study tested whether intrinsic contractile function of human skeletal muscle fibers is impaired in older adults by integrating single fiber contractile mechanics with 3D morphological imaging of the same fibers. We also evaluated the agreement between conventional methods of estimating fiber CSA from 2D images in solution and air with CSA-derived measurements from 3D imaging (Fig. 3). As expected, peak power of the knee extensor muscles was closely associated with the involuntary, electrically evoked contractile properties of the potentiated twitch, but not with voluntary activation, suggesting the age-related decline in peak power is due primarily to factors within the muscle (Wrucke *et al*., 2024). Consistent with our previous work (Sundberg *et al*., 2018a; Teigen *et al*., 2020; Sundberg *et al*., 2025; Teigen *et al*., 2026), absolute P_o_ and peak power were markedly lower in fast MyHC II fibers from older adults but did not differ with age in slow MyHC I fibers. However, when normalized to CSA estimated from 2D images in air or derived from 3D imaging, size-specific force and power of both MyHC I and II fibers either did not differ with age or were greater in older compared with young adults, indicating preserved intrinsic contractile function with aging. In contrast, normalization to CSA estimated from 2D images in solution resulted in lower size-specific power in MyHC I fibers and lower size-specific P_o_ in MyHC II fibers in the older adults (Figs. 6 and 7). Importantly, all 2D methods systematically overestimated 3D-derived fiber CSA, with solution-based estimates demonstrating the greatest bias and poorest agreement with 3D imaging (Fig. 3). These data suggest that fast fiber atrophy is a major contributor to declines in whole muscle force and power in older adults, and that conventional methods to estimate fiber CSA can influence conclusions regarding age-related differences in intrinsic contractile function, providing a probable explanation for the longstanding inconsistencies in the literature.

### Intrinsic contractile function is preserved with aging despite pronounced fast fiber atrophy

A novel aspect of the present study was the development of a custom imaging microchamber system (Fig. 1) that enabled integration of single fiber contractile mechanics with high-resolution 3D morphological measurements of the same fiber segments at a fixed sarcomere length (2.5 μm). This approach eliminated the reliance on assumptions of fiber cross-sectional shape that is inherent to conventional single fiber methods and allowed CSA to be quantified along the length of the fiber (∼635 μm) at a spatial resolution of 0.38 μm^2^. Using this approach, we found that size-specific force and power did not differ between young and older males in either fiber type, indicating that intrinsic contractile function is preserved with aging. Consistent with this interpretation, we also observed no age-related differences in V_max_ or the rate of tension redevelopment (k_tr_), both of which reflect intrinsic contractile kinetics that are independent of fiber size. Furthermore, by directly comparing 3D-derived CSA with estimates obtained from conventional 2D imaging methods, the present study provides insight into the longstanding inconsistencies in the single fiber literature regarding whether intrinsic contractile function is impaired with aging.

As expected, fast MyHC II fibers from older adults exhibited lower CSA (Fig. 4B), absolute force (P_o_) (Fig. 5B), and peak power (Fig. 7B), with no age-related differences observed in slow MyHC I fibers, a finding consistent with our previous work (Sundberg *et al*., 2018a; Teigen *et al*., 2020; Sundberg *et al*., 2025; Teigen *et al*., 2026) and others (Gries *et al*., 2019; Grosicki *et al*., 2021). However, when CSA estimated from 2D images in solution was used for size normalization, size-specific power in MyHC I fibers and size-specific P_o_ in MyHC II fibers were lower in older adults, which would suggest that aging impairs intrinsic contractile function (Figs. 6 and 7). Notably, nearly all studies reporting lower size-specific P_o_ in slow MyHC I or fast MyHC II fibers from older adults also estimated fiber size in solution (Larsson *et al*., 1997; Frontera *et al*., 2000; D’Antona *et al*., 2003; D’Antona *et al*., 2007; Ochala *et al*., 2007; Yu *et al*., 2007; Hvid *et al*., 2013; Lamboley *et al*., 2015; Brocca *et al*., 2017). In contrast, only one study using air-based measurements reported an age-related reduction in size-specific P_o_ in MyHC I fibers, with no difference in MyHC IIa fibers (Harber *et al*., 2012). In the present study, CSA estimated in solution was markedly larger than values obtained from air-based measurements or 3D imaging (Fig. 4), which would systematically reduce size-specific P_o_ and power. Consistent with this observation, Degens and Larsson (2007) reported that size-specific P_o_ based on CSA estimated in solution assuming a circular shape was less than 70% of that when assuming an elliptical shape or when CSA was estimated in air. This is comparable to the present findings, in which size-specific P_o_ and power measured in solution were ∼12-44% lower than values derived from air and 3D measurements across both fiber types and independent of age.

One potential explanation for the larger CSA values observed from 2D images in solution is fiber swelling following chemical permeabilization, which is commonly reported to increase fiber dimensions by ∼20% (Godt & Maughan, 1977; Frontera *et al*., 2000; Ochala *et al*., 2007; Yu *et al*., 2007), although the extent of swelling can vary considerably between individuals and fiber types (Monti *et al*., 2021). However, because both the solution-based CSA estimates from 2D images and the 3D imaging in the present study were obtained under similar conditions (i.e., 2.5 µm sarcomere spacing in relaxing solution), any global swelling would be expected to affect both approaches similarly. Additionally, among the studies that estimated CSA in solution and reported lower size-specific P_o_ in fibers from older adults, only 3 of 9 accounted for fiber swelling (Frontera *et al*., 2000; Ochala *et al*., 2007; Yu *et al*., 2007), suggesting that swelling alone is unlikely to explain the discrepancies in the literature. Instead, our data suggests that the most probable explanation is due to limitations inherent to 2D projection-based measurements and assumptions of fiber geometry. Indeed, fibers from older adults, particularly MyHC II fibers, deviate substantially from simple cylindrical or elliptical geometries (Kirkeby & Garbarsch, 2000; Barnouin *et al*., 2017; Messa *et al*., 2020; Horwath *et al*., 2024; Soendenbroe *et al*., 2024). Consistent with this observation, transferring fibers from solution to air in the present study resulted in a much greater reduction in CSA in fibers from older (26%) compared with young adults (7%), especially in MyHC II fibers. These findings indicate that fibers, particularly those from older adults, deviate markedly from idealized circular or elliptical geometry, and that this deviation is likely contributing to the apparent age-related differences in intrinsic contractile function when fiber CSA is estimated from 2D images in solution.

In contrast to the findings obtained using CSA estimated in solution, air-based measurements and 3D imaging yielded consistent evidence of preserved intrinsic contractile function with aging. However, air-based measurements may also introduce systematic bias, albeit in the opposite direction. Indeed, in the present study, size-specific power was greater in fast MyHC II fibers from older compared with young adults, while 3D imaging indicated no differences (Fig. 7). Notably, most studies reporting higher size-specific P_o_ or peak power in slow MyHC I or fast MyHC II fibers from older adults also estimated fiber size while fibers were suspended in air (Trappe *et al*., 2003; Sundberg *et al*., 2018a; Gries *et al*., 2019; Grosicki *et al*., 2021; Sundberg *et al*., 2025). In contrast, only two studies using solution-based CSA estimates reported greater size-specific P_o_ in MyHC IIa fibers from older adults (Miller *et al*., 2013; Straight *et al*., 2018). Importantly, both studies used a prism to obtain side-view images of the fibers and estimated CSA assuming an elliptical geometry, suggesting that air-based measurements and solution-based measurements incorporating elliptical assumptions using a prism may similarly bias conclusions toward apparent enhancements in intrinsic contractile function with aging. These findings, when interpreted together, provide compelling evidence that methodological differences in CSA estimation, whether performed in solution or air, can bias the conclusions in potentially opposite directions, and contribute to the longstanding inconsistencies in the literature regarding altered intrinsic contractile function with aging.

An important limitation of the present study is that CSA estimates from the 2D images were calculated assuming fibers formed a circular cross-sectional shape, whereas many previous single fiber studies using solution-based CSA estimates have assumed an elliptical geometry (Grosicki *et al*., 2022), which as discussed above may influence measurements of size-specific contractile function. For example, second harmonic generation microscopy of skinned rabbit soleus fibers showed that CSA estimates using circular assumptions resulted in a modestly greater overestimation of real CSA (5.3 ± 25.9%) compared with elliptical assumptions (2.8 ± 6.9%) (Mebrahtu *et al*., 2024). These findings are consistent with earlier work in isolated frog fibers showing that CSA estimates based on a single width measurement are more prone to error than estimates incorporating measurements from multiple axes (Blinks, 1965). However, a recent study in human skeletal muscle fibers reported no age-related differences in size-specific P_o_ when CSA was estimated assuming either circular or elliptical geometry (Kalakoutis *et al*., 2023). Furthermore, a systematic review and meta-analysis demonstrated substantial inter-study variability in size-specific P_o_ of both fiber types from healthy young adults even after accounting for factors such as fiber swelling, sarcomere length, and assumptions of cross-sectional shape (Kalakoutis *et al*., 2021). Collectively, these findings suggest that assumptions regarding fiber geometry likely contribute to discrepancies in the literature regarding size-specific contractile function. However, additional studies directly comparing the methodological approaches are needed to determine the extent to which these inconsistencies reflect differences in measurement techniques versus true biological variability.

Despite the methodological implications with estimating fiber CSA from 2D images, the convergence of findings from the 3D imaging, air-based CSA measurements, and size-independent contractile measurements (k_tr_ and V_max_) provide compelling evidence that intrinsic contractile function is largely preserved with aging, and that the primary age-related alteration at the single fiber level is the markedly lower absolute force and power output of fast fibers (Figs. 5B and 7B).

Given that fast fibers generate ∼0.3-2.5-fold greater size-specific force and power than slow fibers, irrespective of the CSA measurement approach (Figs. 6C and 7C), their selective atrophy would be expected to disproportionately reduce whole muscle force and power relative to the loss of muscle mass. Accordingly, the power output of the fast MyHC II fibers was closely associated with whole muscle power, whereas there was no association with slow MyHC I fiber power (Fig. 2B). When considered alongside the ∼46% lower fast MyHC II proportional area in older adults and the age-related reductions in mass-specific torque (∼22%) and power (∼31%), these findings collectively implicate fast fiber atrophy as an important factor contributing to the reduced whole muscle function with aging. Notably, multiple variables obtained from the involuntary, electrically evoked twitch, but not voluntary activation, were closely associated with knee extensor power output, providing direct evidence that the age-related loss in peak power is driven primarily by factors within the muscle. Future studies focused on identifying the mechanisms contributing to age-related fast fiber atrophy will be important for developing targeted therapeutic strategies to attenuate the loss in muscle strength and power with advancing age.

### Conclusions

This study demonstrates that methods commonly used in single fiber contractile mechanics experiments for estimating CSA from 2D images overestimate values obtained from high-resolution 3D imaging, which has important implications for calculating size-specific force and power. As expected, absolute force and power output were lower in fast fibers from older compared with young adults, whereas no age-related differences were observed in slow fibers. Measurements of size-specific force and power when CSA was estimated in solution were lower in the older compared with young adults but were higher or not different when size was determined from air-based measurements or 3D imaging. We conclude that intrinsic contractile function is preserved in the slow and fast fibers of older males, and that the methods used to estimate fiber size can influence apparent age-related differences in muscle fiber contractile function.

## Data Availability Statement

The data in the present study are available from the corresponding author upon reasonable request.

## Author Contributions

CSZ, LET, and CWS conceived and designed the experiments; CSZ, LET, ID, and CWS performed experiments; CSZ, LET, ID, and YW analyzed the data; CSZ, LET, and CWS interpreted the data; CSZ, LET, ID, and CWS prepared the figures; CSZ, LET, and CWS drafted the manuscript; CSZ, LET, ID, YW, and CWS edited and revised the manuscript; CSZ, LET, ID, YW, and CWS approved the final version of the manuscript. All authors approved the final version of the manuscript and agree to be accountable for all aspects of the work. All persons designated as authors qualify for authorship, and all those who qualify for authorship are listed.

## Acknowledgements

We thank Dan Holbus for assistance with the design and machining of the custom imaging microchamber system, David Wrucke, Jessica James, and Kam Kroner for assistance with the whole-muscle data collection and analysis, and the research participants for volunteering to make this study possible. We are also deeply grateful to Dr. Robert Fitts for gifting his single-fiber contractile mechanics stations to the Sundberg laboratory.

## Appendix (Tables A1 & A2)

**Table A1.**
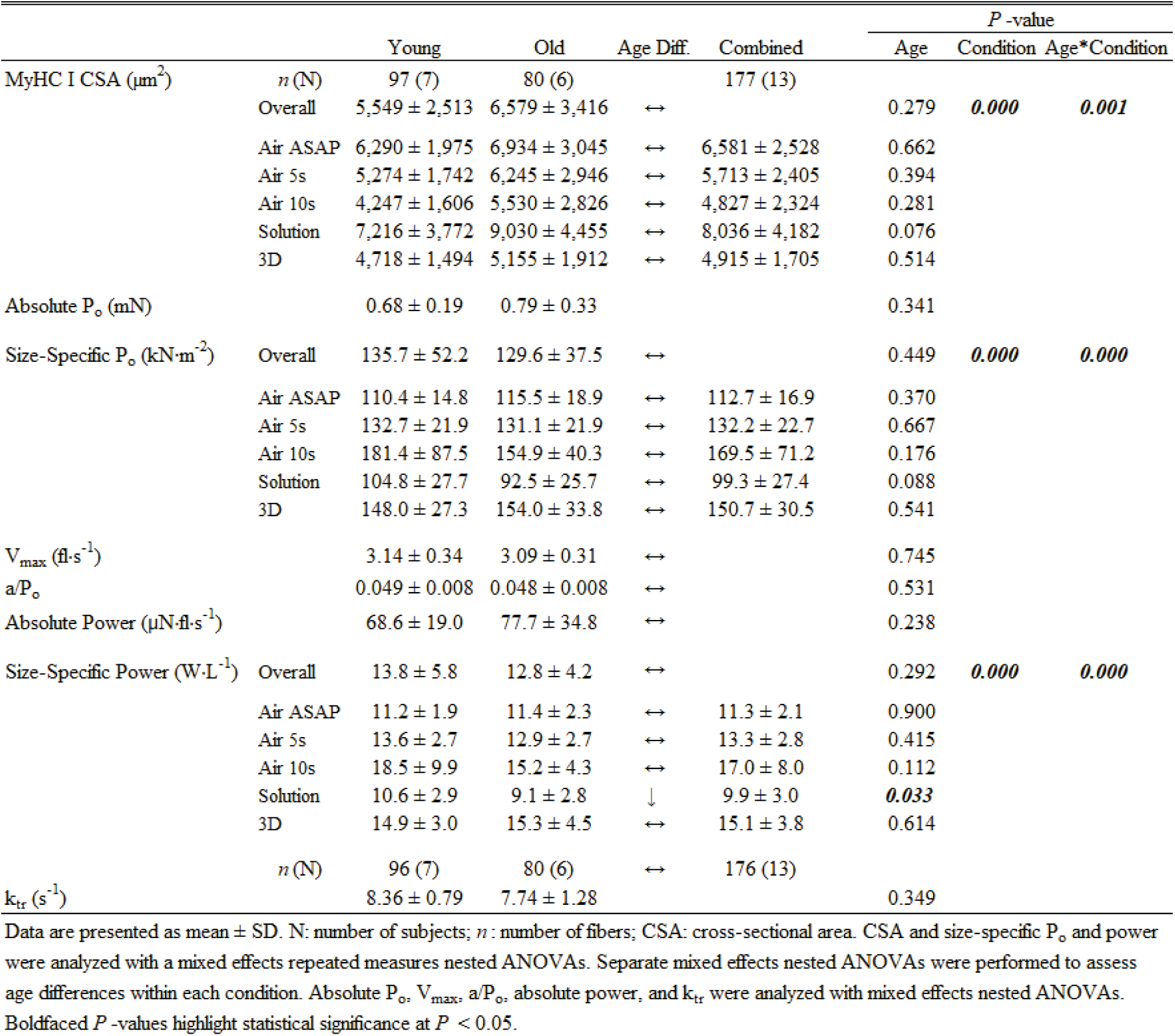
MyHC I single fiber cross-sectional area & contractile mechanics.

**Table A2.** MyHC II single fiber cross-sectional area & contractile mechanics.

|  |  |  |  |  |  | *P*-value |  |  |
| --- | --- | --- | --- | --- | --- | --- | --- | --- |
|  |  | Young | Old | Age Diff. | Combined | Age | Condition | Age*Condition |
| MyHC II CSA ($\mu\text{m}^2$) | *n* (N) | 113 (7) | 87 (6) | | 200 (13) | | | |
| | Overall | 7,841 ± 3,537 | 4,767 ± 2,697 | ↓ | | **0.003** | **0.000** | **0.000** |
| | Air ASAP | 8,902 ± 3,094 | 4,920 ± 1,820 | ↓ | 7,170 ± 3,277 | **0.002** | | |
| | Air 5s | 7,692 ± 2,866 | 4,304 ± 1,670 | ↓ | 6,218 ± 2,943 | **0.004** | | |
| | Air 10s | 6,504 ± 2,697 | 3,626 ± 1,676 | ↓ | 5,252 ± 2,712 | **0.005** | | |
| | Solution | 10,183 ± 4,608 | 7,369 ± 4,088 | ↓ | 8,959 ± 4,597 | **0.034** | | |
| | 3D | 5,922 ± 2,116 | 3,615 ± 1,162 | ↓ | 4,918 ± 2,102 | **0.001** | | |
| Absolute $P_o$ (mN) | | 1.33 ± 0.46 | 0.79 ± 0.28 | ↓ | | **0.003** | | |
| Size-Specific $P_o$ (kN·m<sup>-2</sup>) | Overall | 181.5 ± 47.2 | 185.7 ± 70.9 | ↔ | | 0.692 | **0.000** | **0.000** |
| | Air ASAP | 150.7 ± 21.2 | 161.4 ± 21.9 | ↔ | 155.4 ± 22.1 | 0.177 | | |
| | Air 5s | 176.1 ± 25.8 | 186.8 ± 26.1 | ↔ | 180.7 ± 26.4 | 0.158 | | |
| | Air 10s | 213.0 ± 39.5 | 241.6 ± 104.4 | ↔ | 225.4 ± 76.0 | 0.137 | | |
| | Solution | 140.1 ± 36.7 | 119.3 ± 37.4 | ↓ | 131.1 ± 38.4 | **0.005** | | |
| | 3D | 227.6 ± 36.3 | 219.6 ± 50.2 | ↔ | 224.1 ± 43.0 | 0.667 | | |
| $V_{\max}$ (fl·s<sup>-1</sup>) | | 4.64 ± 0.44 | 4.86 ± 0.54 | ↔ | | 0.056 | | |
| $a/P_o$ | | 0.085 ± 0.020 | 0.095 ± 0.028 | ↔ | | 0.069 | | |
| Absolute Power ($\mu\text{N}\cdot\text{fl}\cdot\text{s}^{-1}$) | | 303.2 ± 134.0 | 192.4 ± 68.0 | ↓ | | **0.020** | | |
| Size-Specific Power (W·L<sup>-1</sup>) | Overall | 41.0 ± 13.4 | 47.7 ± 27.2 | ↑ | | **0.026** | **0.000** | **0.000** |
| | Air ASAP | 34.1 ± 8.6 | 40.8 ± 9.0 | ↑ | 37.0 ± 9.3 | **0.014** | | |
| | Air 5s | 39.8 ± 10.0 | 47.6 ± 13.0 | ↑ | 43.2 ± 12.0 | **0.014** | | |
| | Air 10s | 48.2 ± 13.1 | 64.2 ± 49.2 | ↑ | 55.1 ± 34.8 | **0.015** | | |
| | Solution | 31.6 ± 10.0 | 30.5 ± 12.1 | ↔ | 31.1 ± 11.0 | 0.626 | | |
| | 3D | 51.3 ± 12.6 | 55.3 ± 15.3 | ↔ | 53.1 ± 13.9 | 0.346 | | |
|  | *n* (N) | 110 (7) | 78 (6) |  | 188 (13) |  |  |  |
| $k_{tr}$ (s<sup>-1</sup>) | | 18.16 ± 3.92 | 19.20 ± 3.97 | ↔ | | 0.409 | | |

Data are presented as mean ± SD. N: number of subjects; *n*: number of fibers; CSA: cross-sectional area. CSA and size-specific $P_o$ and power were analyzed with a mixed effects repeated measures nested ANOVAs. Separate mixed effects nested ANOVAs were performed to assess age differences within each condition. Absolute $P_o$ , $V_{\max}$ , $a/P_o$ , absolute power, and $k_{tr}$ were analyzed with mixed effects nested ANOVAs.
Boldfaced *P*-values highlight statistical significance at *P* < 0.05.

## REFERENCES

Alcazar J, Aagaard P, Haddock B, Kamper RS, Hansen SK, Prescott E, Alegre LM, Frandsen U & Suetta C. (2020). Age- and Sex-Specific Changes in Lower-Limb Muscle Power Throughout the Lifespan. The journals of gerontology Series A, Biological sciences and medical sciences 75, 1369–1378.

Alcazar J, Rodriguez-Lopez C, Delecluse C, Thomis M & Van Roie E. (2023). Ten-year longitudinal changes in muscle power, force, and velocity in young, middle-aged, and older adults. Journal of cachexia, sarcopenia and muscle 14, 1019–1032.

Allen JD & Moss RL. (1987). Factors influencing the ascending limb of the sarcomere length-tension relationship in rabbit skinned muscle fibres. J Physiol 390, 119–136.

Araujo CGS, Kunutsor SK, Eijsvogels TMH, Myers J, Laukkanen JA, Hamar D, Niebauer J, Bhattacharjee A, de Souza ESCG, Franca JF & Castro CLB. (2025). Muscle Power Versus Strength as a Predictor of Mortality in Middle-Aged and Older Men and Women. Mayo Clin Proc.

Barnouin Y, McPhee JS, Butler-Browne G, Bosutti A, De Vito G, Jones DA, Narici M, Behin A, Hogrel JY & Degens H. (2017). Coupling between skeletal muscle fiber size and capillarization is maintained during healthy aging. J Cachexia Sarcopenia Muscle 8, 647–659.

Bergstrom J. (1975). Percutaneous needle biopsy of skeletal muscle in physiological and clinical research. Scand J Clin Lab Invest 35, 609–616.

Blinks JR. (1965). Influence of Osmotic Strength on Cross-Section and Volume of Isolated Single Muscle Fibres. J Physiol 177, 42–57.

Brocca L, McPhee JS, Longa E, Canepari M, Seynnes O, De Vito G, Pellegrino MA, Narici M & Bottinelli R. (2017). Structure and function of human muscle fibres and muscle proteome in physically active older men. J Physiol 595, 4823–4844.

Cristea A, Qaisar R, Edlund PK, Lindblad J, Bengtsson E & Larsson L. (2010). Effects of aging and gender on the spatial organization of nuclei in single human skeletal muscle cells. Aging Cell 9, 685–697.

D’Antona G, Pellegrino MA, Adami R, Rossi R, Carlizzi CN, Canepari M, Saltin B & Bottinelli R. (2003). The effect of ageing and immobilization on structure and function of human skeletal muscle fibres. J Physiol 552, 499–511.

D’Antona G, Pellegrino MA, Carlizzi CN & Bottinelli R. (2007). Deterioration of contractile properties of muscle fibres in elderly subjects is modulated by the level of physical activity. Eur J Appl Physiol 100, 603–611.

Degens H & Larsson L. (2007). Application of skinned single muscle fibres to determine myofilament function in ageing and disease. J Musculoskelet Neuronal Interact 7, 56–61.

Delgadillo JD, Sundberg CW, Kwon M & Hunter SK. (2021). Fatigability of the knee extensor muscles during high-load fast and low-load slow resistance exercise in young and older adults. Experimental gerontology 154, 111546.

Fabiato A. (1988). Computer programs for calculating total from specified free or free from specified total ionic concentrations in aqueous solutions containing multiple metals and ligands. Methods in enzymology 157, 378–417.

Fabiato A & Fabiato F. (1979). Calculator programs for computing the composition of the solutions containing multiple metals and ligands used for experiments in skinned muscle cells. Journal de physiologie 75, 463–505.

Ferenczi MA. (1986). Phosphate burst in permeable muscle fibers of the rabbit. Biophys J 50, 471–477.

Frontera WR, Suh D, Krivickas LS, Hughes VA, Goldstein R & Roubenoff R. (2000). Skeletal muscle fiber quality in older men and women. American journal of physiology Cell physiology 279, C611–618.

Godt RE & Maughan DW. (1977). Swelling of Skinned Muscle Fibers of the Frog. Biophysical Journal 19, 103–116.

Gries KJ, Minchev K, Raue U, Grosicki GJ, Begue G, Finch WH, Graham B, Trappe TA & Trappe S. (2019). Single-muscle fiber contractile properties in lifelong aerobic exercising women. Journal of applied physiology (Bethesda, Md: 1985) 127, 1710–1719.

Grosicki GJ, Gries KJ, Minchev K, Raue U, Chambers TL, Begue G, Finch H, Graham B, Trappe TA & Trappe S. (2021). Single muscle fibre contractile characteristics with lifelong endurance exercise. J Physiol 599, 3549–3565.

Grosicki GJ, Zepeda CS & Sundberg CW. (2022). Single muscle fibre contractile function with ageing. J Physiol 600, 5005–5026.

Harber MP, Konopka AR, Undem MK, Hinkley JM, Minchev K, Kaminsky LA, Trappe TA & Trappe S. (2012). Aerobic exercise training induces skeletal muscle hypertrophy and age-dependent adaptations in myofiber function in young and older men. Journal of applied physiology (Bethesda, Md: 1985) 113, 1495–1504.

Hart TL, Swartz AM, Cashin SE & Strath SJ. (2011). How many days of monitoring predict physical activity and sedentary behaviour in older adults? The international journal of behavioral nutrition and physical activity 8, 62.

Hassanlouei H, Sundberg CW, Smith AE, Kuplic A & Hunter SK. (2017). Physical activity modulates corticospinal excitability of the lower limb in young and old adults. Journal of applied physiology (Bethesda, Md: 1985) 123, 364–374.

Herbert RD & Gandevia SC. (1999). Twitch interpolation in human muscles: mechanisms and implications for measurement of voluntary activation. J Neurophysiol 82, 2271–2283.

Hessel AL, Manross KM, Borkowski MM, Rand CD & Nguyen K. (2026). A primer on the methods of skeletal and cardiac muscle mechanics using permeabilized preparations. J Gen Physiol 158.

Hill AV. (1938). The heat of shortening and the dynamic constants of muscle. Proceedings of the Royal Society of London Series B - Biological Sciences 126, 136–195.

Horwath O, Moberg M, Edman S, Philp A & Apro W. (2024). Ageing leads to selective type II myofibre deterioration and denervation independent of reinnervative capacity in human skeletal muscle. Exp Physiol.

Hvid LG, Suetta C, Aagaard P, Kjaer M, Frandsen U & Ortenblad N. (2013). Four days of muscle disuse impairs single fiber contractile function in young and old healthy men. Experimental gerontology 48, 154–161.

Kalakoutis M, Di Giulio I, Douiri A, Ochala J, Harridge SDR & Woledge RC. (2021). Methodological considerations in measuring specific force in human single skinned muscle fibres. Acta physiologica (Oxford, England) 233, e13719.

Kalakoutis M, Pollock RD, Lazarus NR, Atkinson RA, George M, Berber O, Woledge RC, Ochala J & Harridge SDR. (2023). Revisiting specific force loss in human permeabilised single skeletal muscle fibres obtained from older individuals. Am J Physiol Cell Physiol.

Kirkeby S & Garbarsch C. (2000). Aging affects different human muscles in various ways. An image analysis of the histomorphometric characteristics of fiber types in human masseter and vastus lateralis muscles from young adults and the very old. Histol Histopathol 15, 61–71.

Korhonen MT, Cristea A, Alen M, Hakkinen K, Sipila S, Mero A, Viitasalo JT, Larsson L & Suominen H. (2006). Aging, muscle fiber type, and contractile function in sprint-trained athletes. Journal of applied physiology (Bethesda, Md: 1985) 101, 906–917.

Lamboley CR, Wyckelsma VL, Dutka TL, McKenna MJ, Murphy RM & Lamb GD. (2015). Contractile properties and sarcoplasmic reticulum calcium content in type I and type II skeletal muscle fibres in active aged humans. J Physiol 593, 2499–2514.

Larsson L, Li X & Frontera WR. (1997). Effects of aging on shortening velocity and myosin isoform composition in single human skeletal muscle cells. The American journal of physiology 272, C638–649.

Lexell J, Taylor CC & Sjöström M. (1988). What is the cause of the ageing atrophy? Journal of the Neurological Sciences 84, 275–294.

Liu JX, Hoglund AS, Karlsson P, Lindblad J, Qaisar R, Aare S, Bengtsson E & Larsson L. (2009). Myonuclear domain size and myosin isoform expression in muscle fibres from mammals representing a 100,000-fold difference in body size. Exp Physiol 94, 117–129.

Maden-Wilkinson TM, Degens H, Jones DA & McPhee JS. (2013). Comparison of MRI and DXA to measure muscle size and age-related atrophy in thigh muscles. Journal of musculoskeletal & neuronal interactions 13, 320–328.

Malakoutian M, Theret M, Yamamoto S, Dehghan-Hamani I, Lee M, Street J, Rossi F, Brown SHM & Oxland TR. (2021). Larger muscle fibers and fiber bundles manifest smaller elastic modulus in paraspinal muscles of rats and humans. Sci Rep 11, 18565.

Mebrahtu A, Smith IC, Liu S, Abusara Z, Leonard TR, Joumaa V & Herzog W. (2024). Reconsidering assumptions in the analysis of muscle fibre cross-sectional area. J Exp Biol 227.

Messa GAM, Piasecki M, Rittweger J, McPhee JS, Koltai E, Radak Z, Simunic B, Heinonen A, Suominen H, Korhonen MT & Degens H. (2020). Absence of an aging-related increase in fiber type grouping in athletes and non-athletes. Scand J Med Sci Sports 30, 2057–2069.

Metzger JM, Greaser ML & Moss RL. (1989). Variations in cross-bridge attachment rate and tension with phosphorylation of myosin in mammalian skinned skeletal muscle fibers. Implications for twitch potentiation in intact muscle. The Journal of general physiology 93, 855–883.

Metzger JM & Moss RL. (1987). Greater hydrogen ion-induced depression of tension and velocity in skinned single fibres of rat fast than slow muscles. J Physiol 393, 727–742.

Miller MS, Bedrin NG, Callahan DM, Previs MJ, Jennings ME, 2nd, Ades PA, Maughan DW, Palmer BM & Toth MJ. (2013). Age-related slowing of myosin actin cross-bridge kinetics is sex specific and predicts decrements in whole skeletal muscle performance in humans. Journal of applied physiology (Bethesda, Md: 1985) 115, 1004–1014.

Mitchell WK, Williams J, Atherton P, Larvin M, Lund J & Narici M. (2012). Sarcopenia, dynapenia, and the impact of advancing age on human skeletal muscle size and strength; a quantitative review. Frontiers in physiology 3, 260.

Monti E, Toniolo L, Marcucci L, Bondì M, Martellato I, Šimunič B, Toninello P, Franchi MV, Narici MV & Reggiani C. (2021). Are muscle fibres of body builders intrinsically weaker? A comparison with single fibres of aged-matched controls. Acta physiologica (Oxford, England) 231, e13557.

Moss RL. (1979). Sarcomere length-tension relations of frog skinned muscle fibres during calcium activation at short lengths. J Physiol 292, 177–192.

Nilwik R, Snijders T, Leenders M, Groen BB, van Kranenburg J, Verdijk LB & van Loon LJ. (2013). The decline in skeletal muscle mass with aging is mainly attributed to a reduction in type II muscle fiber size. Exp Gerontol 48, 492–498.

Ochala J, Frontera WR, Dorer DJ, Van Hoecke J & Krivickas LS. (2007). Single skeletal muscle fiber elastic and contractile characteristics in young and older men. The journals of gerontology Series A, Biological sciences and medical sciences 62, 375–381.

Raue U, Slivka D, Minchev K & Trappe S. (2009). Improvements in whole muscle and myocellular function are limited with high-intensity resistance training in octogenarian women. Journal of applied physiology (Bethesda, Md: 1985) 106, 1611–1617.

Reid KF & Fielding RA. (2012). Skeletal muscle power: a critical determinant of physical functioning in older adults. Exerc Sport Sci Rev 40, 4–12.

Smith IC & Herzog W. (2023). Assumptions about the cross-sectional shape of skinned muscle fibers can distort the relationship between muscle force and cross-sectional area. J Appl Physiol (1985) 135, 1036–1040.

Soendenbroe C, Karlsen A, Svensson RB, Kjaer M, Andersen JL & Mackey AL. (2024). Marked irregular myofiber shape is a hallmark of human skeletal muscle ageing and is reversed by heavy resistance training. J Cachexia Sarcopenia Muscle 15, 306–318.

Straight CR, Ades PA, Toth MJ & Miller MS. (2018). Age-related reduction in single muscle fiber calcium sensitivity is associated with decreased muscle power in men and women. Experimental gerontology 102, 84–92.

Sundberg CW, Hunter SK, Trappe SW, Smith CS & Fitts RH. (2018a). Effects of elevated H(+) and Pi on the contractile mechanics of skeletal muscle fibres from young and old men: implications for muscle fatigue in humans. J Physiol 596, 3993–4015.

Sundberg CW, Kuplic A, Hassanlouei H & Hunter SK. (2018b). Mechanisms for the age-related increase in fatigability of the knee extensors in old and very old adults. Journal of applied physiology (Bethesda, Md: 1985) 125, 146–158.

Sundberg CW, Teigen LE, Hunter SK & Fitts RH. (2025). Cumulative effects of H(+) and P(i) on force and power of skeletal muscle fibres from young and older adults. J Physiol 603, 187–209.

Teigen LE, Sundberg CW, Kelly LJ, Hunter SK & Fitts RH. (2020). Ca(2+) dependency of limb muscle fiber contractile mechanics in young and older adults. American journal of physiology Cell physiology 318, C1238–c1251.

Teigen LE, Zepeda CS, Dobrzycki I, Hunter SK, Fitts RH & Sundberg CW. (2026). Effects of acidosis and inorganic phosphate on Ca(2+) sensitivity of young and older adult skeletal muscle fibers. Am J Physiol Cell Physiol 330, C224–C237.

Trappe S, Gallagher P, Harber M, Carrithers J, Fluckey J & Trappe T. (2003). Single muscle fibre contractile properties in young and old men and women. J Physiol 552, 47–58.

Wen Y, Murach KA, Vechetti IJ, Jr., Fry CS, Vickery C, Peterson CA, McCarthy JJ & Campbell KS. (2018). MyoVision: software for automated high-content analysis of skeletal muscle immunohistochemistry. J Appl Physiol (1985) 124, 40–51.

Widrick JJ, Trappe SW, Costill DL & Fitts RH. (1996). Force-velocity and force-power properties of single muscle fibers from elite master runners and sedentary men. The American journal of physiology 271, C676–683.

Wrucke DJ, Colosio M, James JJ, Hunter SK & Sundberg CW. (2026). Initial contractions of a single-load fatiguing exercise provide a valid assessment of age-related differences in peak power. J Appl Physiol (1985).

Wrucke DJ, Kuplic A, Adam MD, Hunter SK & Sundberg CW. (2024). Neural and muscular contributions to the age-related differences in peak power of the knee extensors in men and women. J Appl Physiol (1985) 137, 1021–1040.

Yu F, Hedström M, Cristea A, Dalén N & Larsson L. (2007). Effects of ageing and gender on contractile properties in human skeletal muscle and single fibres. Acta physiologica (Oxford, England) 190, 229–241.

